# TSST-1 promotes colonization of *Staphylococcus aureus* within the vaginal tract by activation of CD8^+^ T cells

**DOI:** 10.1101/2024.10.04.616698

**Authors:** Karine Dufresne, Kait F. Al, Heather C. Craig, Charlotte E.M. Coleman, Katherine J. Kasper, Jeremy P. Burton, John K. McCormick

**Affiliations:** Department of Microbiology and Immunology, Schulich School of Medicine and Dentistry, University of Western Ontario, London (ON), Canada; Canadian Centre for Human Microbiome and Probiotics Research, Lawson Health Research Institute, London, Ontario, Canada

**Keywords:** *Staphylococcus aureus*, superantigen, TSST-1, T cells, vaginal environment

## Abstract

Toxic shock syndrome toxin-1 (TSST-1) is a superantigen produced by *Staphylococcus aureus* and is the determinant of menstrual toxic shock syndrome (mTSS); however, the impact of TSST-1 on the vaginal environment beyond mTSS is not understood. Herein, we assessed how TSST-1 affects vaginal colonization by *S. aureus*, host inflammatory responses, and changes in microbial communities within the murine vagina. We demonstrated that TSST-1 induced a CD8^+^ T cell-dependent inflammatory response by 24 hours that correlated with an increased bacteria burden within the vaginal tract. This increase was due to superantigen-dependent T cell activation that triggered a change in microbial composition within the vaginal tract. Altogether, this study demonstrates that within the vaginal tract, TSST-1 modulates the vaginal microbiota to favor the survival of *S. aureus* in the absence of mTSS.

**Importance:** Toxic shock syndrome toxin-1 (TSST-1) is a superantigen toxin produced from *Staphylococcus aureus* that causes the menstrual form of toxic shock syndrome. This research demonstrates that TSST-1 also has a wider function within the vaginal tract than previously expected. We show that TSST-1, by activating CD8^+^ T cells, induces an inflammatory environment that modifies the vaginal microbiota to favor colonization by *S. aureus*. These are important findings as *S. aureus* can colonize the human vaginal tract efficiently and subsequently trigger dysbiosis within the microbial communities leading to several adverse outcomes such as decreased fertility, increased risks for sexually transmitted diseases and issues related to pregnancy and birth.

## Introduction

Approximately one-third of the human population is thought to be chronically colonized by *Staphylococcus aureus* either on the skin or other mucosal surfaces including the nasal and vaginal tracts (1). However, colonization by *S. aureus* can become harmful when natural host defenses are weakened leading to numerous diseases ranging from superficial skin infections to life-threatening invasive conditions including bacteremia, endocarditis and pneumonia (2).

Apart from a broad range of infections, *S. aureus* can also cause specific toxin-mediated diseases including the toxic shock syndrome (TSS) which is triggered by the bacterial superantigens (SAgs) (3). SAgs are secreted exotoxins that short-circuit the interaction between major histocompatibility class II (MHC-II) molecules and T cell receptors (TCRs), which may lead to uncontrolled T cell activation, excessive cytokine production and systemic inflammation known as TSS (4, 5). Staphylococcal TSS can be distinguished into two subgroups where the non-menstrual form can occur from essentially any *S. aureus* infection and can be caused by different SAgs; however, the menstrual form of TSS (mTSS) is usually related to the use of menstrual hygiene products such as tampons and occurs in women who are vaginally colonized by *S. aureus* that specifically produce the toxic shock syndrome toxin-1 (TSST-1) (3). Nevertheless, little is known about how TSST-1 functions within the *S. aureus* life cycle.

*S. aureus* is a well-adapted human colonizer, yet few studies have assessed the determinants of colonization within the vaginal tract. Gajer et al (2012) characterized the human vaginal microbiota during a time course of 16 weeks, where the majority of the 32 women tested harbored staphylococcal species at least once during the sampling period (6). Additionally, Chiaruzzi et al (2020) identified the presence of *S. aureus* on tampons of 27% of healthy women (7). These two studies suggest that the presence of *S. aureus* in the vagina may be underestimated and may depend on the menstrual cycle and other dynamics of the vaginal tract. Jacquemond et al (2018) proposed that the presence of *S. aureus* alters the composition of the microbial community suggesting a more important involvement of the bacterium in the vaginal environment than previously defined (8). Finally, Deng et al (2019) evaluated key determinants of methicillin-resistant *S. aureus* (MRSA) within the vaginal tract and identified that iron acquisition systems and fibrinogen binding proteins were important for successful colonization (9). As TSST-1 is the major determinant of mTSS, one key question that remains unanswered is the role that this superantigen plays in earlier steps of *S. aureus* colonization and pathogenesis within the vaginal tract.

In this study, we proposed to decipher the role of TSST-1 in vaginal colonization by *S. aureus* in a BALB/c mouse model that elicits a TSST-1-driven immune response. We hypothesized that the SAg TSST-1 would be important to enable successful colonization by modulation of the immune response in the vaginal tract conferring an advantage to the bacterium against the endogenous microbiota. Understanding the colonization and activity of *S. aureus* in the female reproductive tract should help lead to strategies that reduce the probability of mTSS and other inflammatory diseases such as staphylococcal-associated aerobic vaginitis (AV).

## Results

### TSST-1 increases *S. aureus* burden in the vaginal tract during diestrus

To study the function of TSST-1 in vaginal colonization, we utilized *S. aureus* MN8, a TSST-1^+^ clinical isolate obtained from a mTSS patient in 1980 (10) (**Table 1**), and generated an in-frame, markerless deletion in the *tst* gene. In addition, we generated a complementation plasmid with the *tst* gene cloned into pCM29 (replacing the vector encoded *gfp* gene). Both the wild-type, the MN8 Δ*tst* and the *tst* complemented strains demonstrated similar growth in tryptic soy broth (TSB) (**Fig 1A**), and although TSST-1 has also been characterized as a repressive gene regulator (11), we saw no evidence of altered protein profiles in the MN8 Δ*tst* strain except for the lack of TSST-1, noting that the protein band visible at ∼25 kDa in the two strains containing the vector corresponds to GFP encoded within pCM29 (**Fig 1B**).

**FIG 1.**
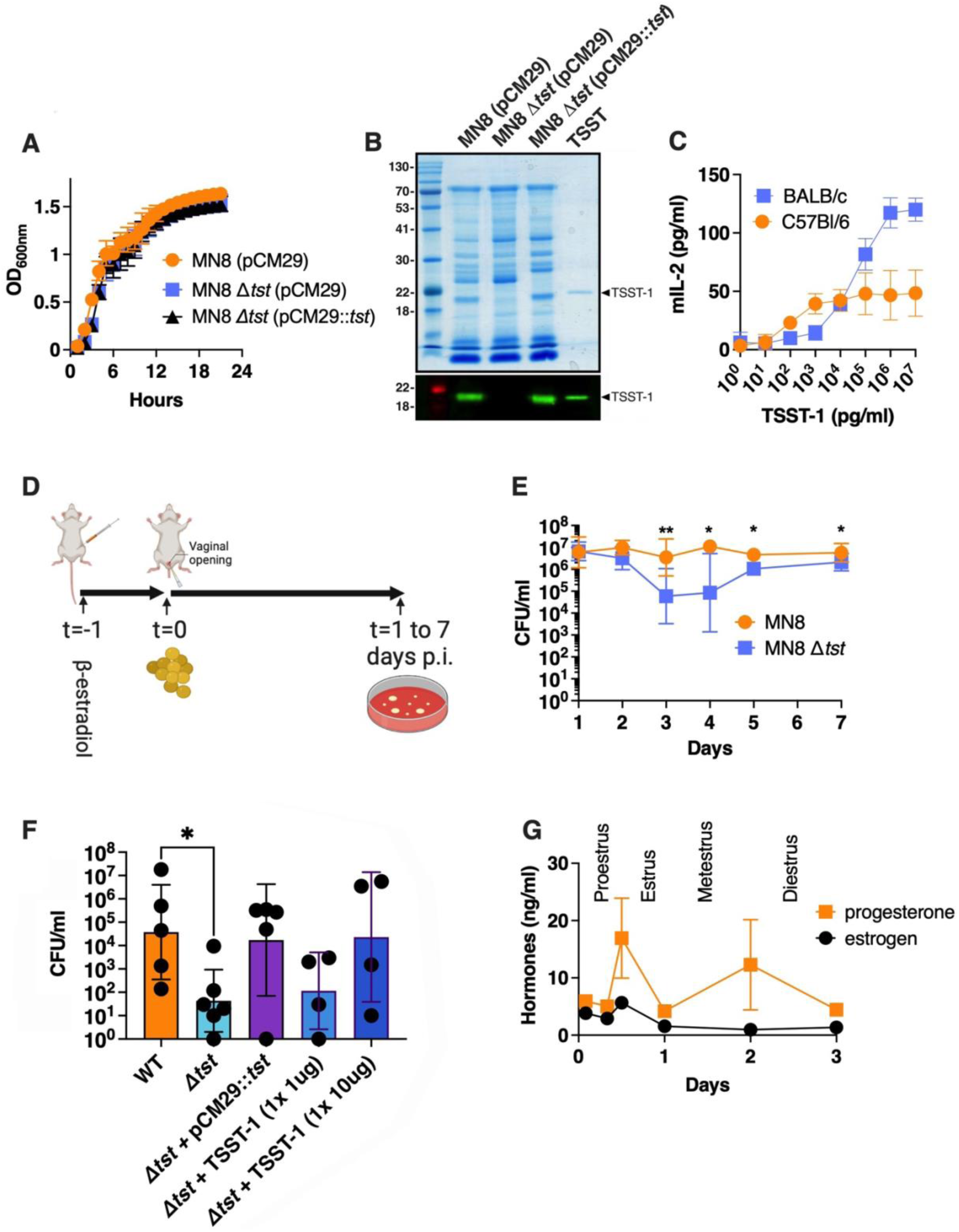
TSST-1 provides an advantage for colonization of the mouse vagina during diestrus. **(A)** Growth of indicated *S. aureus* strains in TSB over 24 hours. Results are presented as the OD measured at 600nm. **(B)** Exoprotein profiles (top panel) and anti-TSST-1 Western blot (bottom panel) for the indicated strains. Concentrated supernatants from the indicated strains grown in TSB medium for 18 h were loaded onto 12% SDS-PAGE gels. Molecular mass markers were loaded on the left and labeled in kilodaltons. Purified recombinant TSST-1 was loaded on the right and is indicated by the solid arrowhead. **(C)** The susceptibility of T cells to TSST-1 from conventional mice was tested by exposing splenocytes from either BALB/c or C57Bl/6 mice with titrating concentrations of recombinant TSST-1. T cell activation was assessed by the production of murine IL-2. Results are presented as the mean of at least 4 biological replicates ± SD. **(D)** Timeline of the vaginal colonization model. The estrous cycle of the animals was synchronized by injecting intraperitoneally 0.5mg/ml of β-estradiol at day 0. **(E)** Wild-type *S. aureus* MN8 or the MN8 Δ*tst* mutant were intravaginally inoculating a dose of ∼10^6^ bacterial CFUs and the bacterial burden within the vaginal tract was assessed from 1 day to 7 days after. Bacterial CFUs were determined observed plating vaginal homogenates on MSA supplemented with chloramphenicol. Data are represented as the geometric mean ± SD. Each dot represents an individual mouse. **(F)** Vaginal colonization was determined at day 3 with wild-type *S. aureus* MN8 or the MN8 *Δtst* strain and complemented with either pCM29::*tst* or with purified TSST-1 as indicated. Results are represented as the geometric mean of at least 4 biological replicates ± SD. **(G)** Hormone levels within the vaginal homogenates were assessed at various timepoints and presented as the mean of at least 4 replicates ± SEM.

**Table 1.** Bacterial strains used in this study.

| Strain | Description | Source |
| --- | --- | --- |
| ***S. aureus*** |  |  |
| MN8 | Prototypic menstrual TSS strain, *tst*<sup>+</sup> | (48) |
| MN8 Δ*tst* | MN8 with deletion of *tst* gene | This study |
| MN8 (pCM29) | MN8 containing pCM29 | This study |
| MN8 Δ*tst* (pCM29) | MN8 Δ*tst* containing pCM29 | This study |
| MN8 Δ*tst* (pCM29::*tst*) | MN8 Δ*tst* containing pCM29::*tst* | This study |
| ***E. coli*** |  |  |
| XL1 Blue | General cloning strain | Stratagene |
| SA30B | DNA methylation strain | (39) |
| BL21(DE3) | Protein expression strain | New England Biolabs |
| BL21(DE3) (pET28a::*tst*) | BL21(DE3) containing the pET28a::*tst* | (17) |
| BL21(DE3) (pET28a::*tst*<sub>S72A</sub>) | BL21(DE3) containing the pET28a::*tst*<sub>S72A</sub> | (17) |

In order to assess the role of the SAg TSST-1 in vaginal colonization, a murine model must be susceptible to toxin activity as in general, mouse MHC-II molecules function poorly with most bacterial superantigens (12–14). To evaluate this, isolated spleen cells from different conventional mouse strains were exposed to purified recombinant TSST-1 (**Fig 1B**) to determine an appropriate mouse strain for this study. Compared to conventional C57BL/6 mice, BALB/c mice demonstrated increased activation of T cells as measured by increased mouse IL-2 levels at higher doses of TSST-1 (**Fig 1C**). These data indicate that BALB/c lymphocytes are susceptible to TSST-1 activity and BALB/c female mice were used for the design of the vaginal colonization model.

Next, the estrous cycle of mice was synchronized with β-estradiol and *S. aureus* MN8, or the MN8 *Δtst* mutant, were inoculated intravaginally (∼2.5 × 10^6^ CFUs) and mice were sacrificed at multiple time points to assess the bacterial burden within the vaginal tract (**Fig 1D**). The recovered cell counts of both strains was similar at the early stages of colonization, however, by day 3 post-inoculation, the MN8 Δ*tst* strain demonstrated ∼100-fold decreased colony-forming units (CFUs) compared to the wild-type strain (**Fig 1E**). This difference was also observed at later timepoints until day 7 post-inoculation where the Δ*tst* strain reached a bacterial burden that was similar to wildtype (**Fig 1E**). We then assessed the ability of the TSST-1 complemented strain (MN8 Δ*tst*+pCM29::*tst*) to colonize the mice and found that this strain was similar to wild-type MN8, demonstrating there was no polar phenotype due to the Δ*tst* deletion (**Fig 1F**). Moreover, an exogenous dose of purified TSST-1 administered at 10μg, but not at 1μg, added at the inoculation of MN8 Δ*tst* within our model was also able to complement the bacterial burden of MN8 Δ*tst* that was similar to wild-type MN8 levels (**Fig 1F**). These data indicate that *S. aureus* MN8 has a colonization advantage within the vaginal environment compared to the MN8 *Δtst* mutant, and this was due to the production of the TSST-1 superantigen.

*S. aureus* persistence can be influenced by the estrous cycles and to evaluate potential influences of estrogen and progesterone, these hormonal levels were assessed from 2 hours to 72 hours post-inoculation in vaginal homogenates, when differences in bacterial burden between wild-type MN8 and MN8 *Δtst* was observed (**Fig 1G**). Hormonal levels were similar between mice inoculated with wildtype MN8 or MN8 *Δtst* and were plotted together (**Fig 1G**). During the period assessed, 2 peaks of progesterone and 1 peak of estrogen were observed, representing the entire estrous cycle (15). The synchronized peaks of both estrogen and progesterone represent the period in between proestrus and estrus, and metestrus starts with the decrease of both hormones before the second peak of progesterone to switch into diestrus following the second peak of progesterone (**Fig 1G**) (15). Overall, these data demonstrate that mice reach the diestrus stage immediately before the observed colonization difference between *S. aureus* MN8 and the MN8 *Δtst* mutant.

### TSST-1 provides a colonization advantage for *S. aureus* against the vaginal microbiota

We hypothesized that the production of TSST-1 may function to provide a colonization advantage to *S. aureus* in competition with the vaginal microbiota through reduced colonization resistance. To evaluate if the microbiota contributed to *S. aureus* burden, mice received fresh water with or without chloramphenicol *ad libitum*, and were intravaginally inoculated with either wild-type *S. aureus* MN8 or the MN8 *Δtst* mutant. The *S. aureus* strains contained the stable pCM29 plasmid to provide chloramphenicol resistance to the antibiotic. The inclusion of chloramphenicol in the water did not alter wild-type *S. aureus* MN8 numbers (**Fig 2A, orange bars**), but conversely, mice colonized with the MN8 Δ*tst* mutant demonstrated significantly decreased numbers only in mice that did not receive chloramphenicol (**Fig 2A, blue bars**). In separate experiments, the culturable microbiota load was assessed from vaginal homogenates by culturing the samples on various media including Man Rogosa and Sharpe agar (MRS), *Enterococcus* Selective medium, 5% sheep blood TSB, and Mannitol salt agar (MSA). Both blood and MSA media without chloramphenicol showed similar numbers of recovered wild-type *S. aureus* MN8 compared with chloramphenicol in the media (data not shown), whereas presumed *Lactobacillus* and *Enterococcus* spp. were relatively low but detectable in the absence of chloramphenicol, but were not detectable from mice that received chloramphenicol containing water (**Fig S1A & S1B**), demonstrating that the chloramphenicol depletion protocol selected against known predominant groups present in the murine vagina (16). These data suggest that TSST-1 may contribute to competition with bacterial residents of the vaginal microbiota.

**FIG 2.**
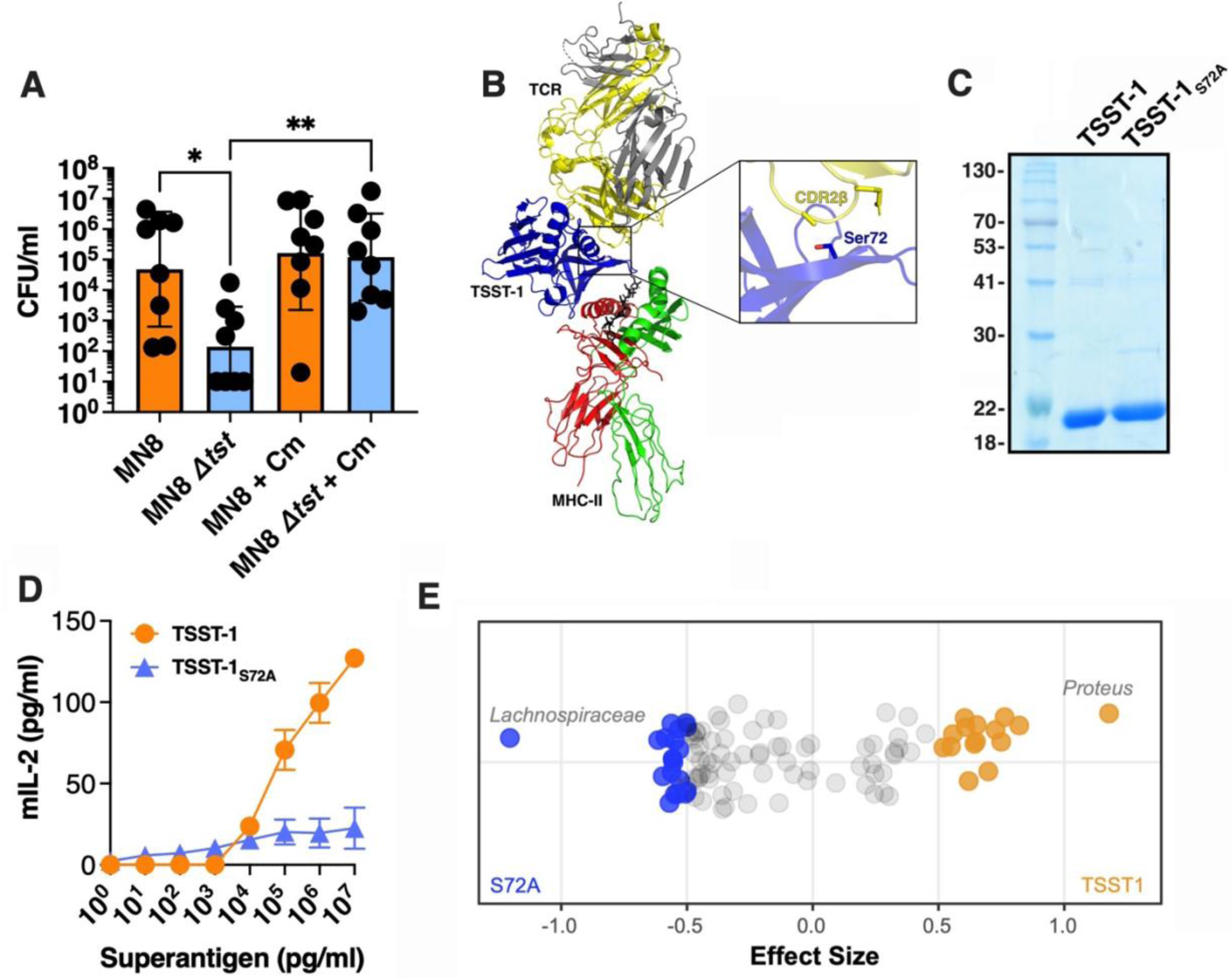
TSST-1 provides a colonization advantage for *S. aureus* against the vaginal microbiota. **(A)** Mice were supplied with water ad libitum with or without chloramphenicol and colonized vaginal inoculated with ∼10^6^ wild-type *S. aureus* MN8 or the MN8 Δ*tst* mutant. Mice were sacrificed at day 3 and data are presented as the geometric mean ± SD (*, p ≤ 0.05; **, p < 0.01). **(B)** Ribbon diagram model of TSST-1 (blue) in complex with the TCR (α-chain, grey; β-chain, yellow) and MHC class II (α-chain, red; β-chain, green). Inset image shows the location of the TSST-1 Ser^72^ amino acid interacting with the CDR2β loop of the TCR β-chain. **(C)** Recombinant TSST-1 and TSST-1_S72A_ visualized on a Coomassie stained 12% SDS-PAGE. **(D)** Activation profile of titrating doses of TSST-1 and TSST-1_S72A_ exposed BALB/c splenocytes. Murine IL-2 production was assessed by ELISA after 18h. The results are presented as the mean of at 3 biological replicates ± SD. **(E)** Microbial taxa were differentially abundant between treatment groups, indicating toxin-dependent microbiota alteration. Each point represents a single taxonomic sequence variant, plotted horizontally according to its effect size of relative enrichment. Taxa with an effect size > |0.5| were deemed statistically significant and are colored by cohort of relative enrichment: orange and blue taxa had significantly higher relative abundance in mice vaginally inoculated with 10 µg of TSST-1 or TSST-1_S72A_, respectively. The genera of the most significantly differential taxa are labelled (see **Table S2** for details on differential taxa).

Next, we sought to determine whether the microbiota composition was affected specifically by the presence of TSST-1. To test this, mice were treated vaginally with either fully active recombinant TSST-1, or with an inactive TSST-1_S72A_ mutant protein used as a negative control. Residue Ser^72^ in TSST-1 is a critical residue important for interaction with the CDR2 loops of the TCR β-chain, and mutation of Ser^72^ to alanine abolishes the ability of TSST-1_S72A_ to activate human T cells (17) (**Fig 2B**). We expressed and purified both recombinant TSST-1 proteins (**Fig 2C**) and tested these proteins for superantigenic activity on mouse spleen cells using mouse IL-2 as an activation readout. Wild-type TSST-1 produced a dose-dependent IL-2 response while the TSST-1 _S72A_ did not induce significant IL-2 production (**Fig 2D**).

Next, 10 μg of either wildtype TSST-1 or TSST-1_S72A_ was administered intravaginally and the microbiota composition was determined through 16S rRNA gene sequencing (17). This removed the effects of external *S. aureus* abundance as previous reports demonstrated that *S. aureus* can represent a dominating resident in murine vaginal microbiota (18). Mice were sacrificed three days post-inoculation and DNA was extracted from the vaginal tissues. Following nucleic acid extraction, three samples from the vaginal tract inoculated with TSST-1_S72A_, and six samples with wild-type TSST-1, were suitable for further analysis as these samples reached a read count of greater than 1000 during the 16S sequencing (**Table S1**). From this experiment, we found substantial inter-mouse variability in the vaginal microbiota but this corroborating previous literature with abundant taxa including *S. aureus*, *Proteus* spp., *Corynebacterium* spp., and *Ligilactobacillus* spp. (especially *L. animalis* and *L. murinus*) (18) (**Table S2**). However, mice intravaginally inoculated with wild-type TSST-1 were markedly enriched in the relative abundance of *Proteus* spp. and conversely decreased in relative abundance of *Lachnospiraceae* spp. (**Fig 2E**). Moreover, the presence of TSST-1 within the vaginal tract decreased other Gram-positive bacteria relatively from the Bacillota (*Lachnospiraceae* spp.) and Actinomycetota phyla (*Lawsonella* sp.), while enriching those in the phylum Pseudomonadota (*Proteus spp.* And *Afipia* sp.), and other Bacillota (*Enterococcus* sp. And *Ligilactobacillus* sp.) (**Table S2**). Altogether, functional TSST-1 resulted in an altered vaginal niche by shifting the compositional abundance of certain resident species of the microbiota.

### Pro-inflammatory responses are enhanced in the vaginal tract colonized with TSST-1 producing *S. aureus*

As bacterial burdens were ∼100-fold higher with wildtype *S. aureus* MN8 compared to the MN8 Δ*tst* mutant by 72 hours post inoculation (**Fig 1E**), and since TSST-1 functions to activate both CD4^+^ and CD8^+^ T cells (19), we next assessed changes in T cell populations within the vaginal tract. Leukocytes were first discriminated from the other vaginal cells as CD45^+^, we eliminated CD19^+^ (B cells) and F4/80^+^ (macrophages) cells, and further identified T cells as CD4^+^ or CD8^+^ (**Fig S2**). There were no significant differences in the percentages of vaginal CD45^+^ populations in mice (**Fig 3A**), nor were there detectable differences in the percentage of CD4^+^ or CD8^+^ T cells (**Fig 3B and 3C**). However, the presence of leukocytes during diestrus in combination with the presence of TSST-1^+^ *S. aureus* could change the inflammatory profile within the vaginal tract and to assess this, concentrations of inflammatory cytokines and chemokines were measured from vaginal homogenates colonized with either *S. aureus* strain. At 24 hours post-inoculation, we noted an enhancement of pro-inflammatory cytokine responses from mice inoculated with wildtype *S. aureus* MN8 compared to the MN8 Δ*tst* mutant (**Fig 3D**), with notably significant increases in IL-1β and MIP-1β (**Fig S3**). However, by 72 hours, the inflammatory response was more similar between the two strains. Although bacteria we not found in either the spleens or livers of vaginally colonized mice (data not shown), indicating that this is not an invasive infection model, we also assessed cytokine responses in serum samples at 24 and 72 hours. Taken together, the cytokine profiles demonstrate differential pro-inflammatory signals primarily within the vaginal tract in the presence of TSST-1 producing *S. aureus*, whereas the response in the bloodstream predominantly occurred at 24 hours and decreased afterwards suggesting a more restricted superantigen-dependent response within the vaginal mucosa (**Fig 3D and Fig S3)**. These data indicate that although there were no differences in the percentages of the T cell subsets, TSST-1 producing *S. aureus* was able to enhance a cytokine driven inflammatory environment within that vaginal tract.

**FIG 3.**
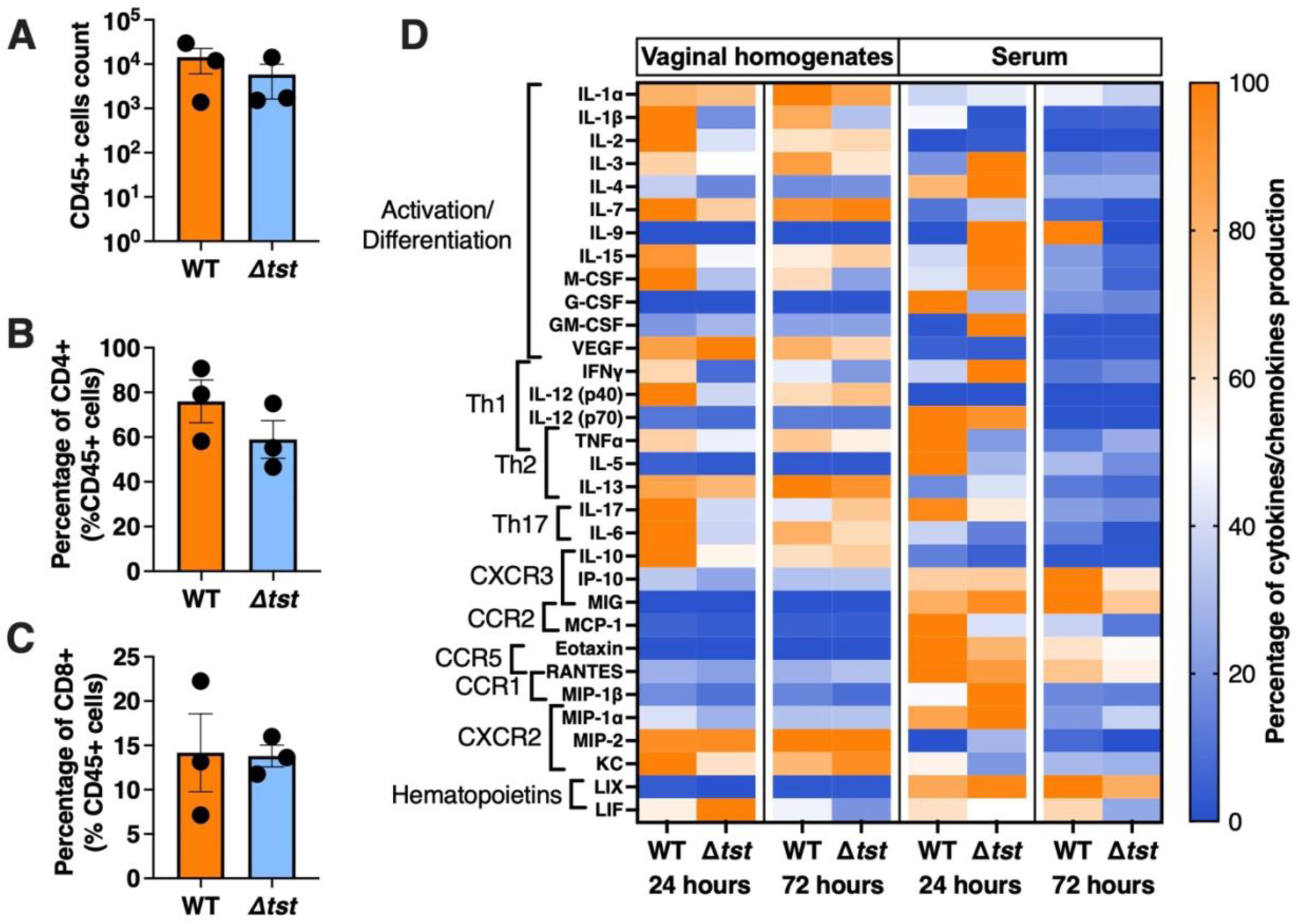
TSST-1^+^ *S. aureus* induced an inflammatory signature within the vaginal tract. **(A-C)** Three days after intra-vaginal inoculation with ∼10^6^ wildtype *S. aureus* MN8 or the Δ*tst* mutant, the number of total immune cells (CD45^+^) and percentages of T cells populations (CD4^+^, CD8^+^) were determined through flow cytometry. Gating strategies are shown in Figure S1. **(D)** Vaginal homogenates and serum from mice vaginally colonized with *S. aureus* MN8 or MN8 *Δtst* were assessed at 24 and 72 hours for inflammatory cytokines and chemokines. The results are represented as the percentage of the averaged production of a specific cytokine compared to its highest concentration set at 100%. Corresponding quantitative data and statistical analyses are shown in **Figure S1**.

### *S. aureus* requires TSST-1 and CD8^+^ T cells to promote colonization fitness within the vaginal tract

Next, to test the functional importance of CD4^+^ and CD8^+^ T cells on *S. aureus* vaginal colonization, we depleted both cell types from the mice both individually and in combination and assessed bacterial burden of wild-type *S. aureus* MN8 colonized mice at 3 days post inoculation (**Fig 4A**). No differences in CFUs were observed between the isotype-treated control animals and mice depleted for CD4^+^ T cells; however, CD8^+^ T cell-depleted animals showed a significant decrease in bacterial by *S. aureus* MN8 (**Fig 4B**). Although CD4^+^/CD8^+^-depleted mice did not show a significant decrease in bacterial burden, a decreased trend was evident (**Fig 4B**).

**FIG 4.**
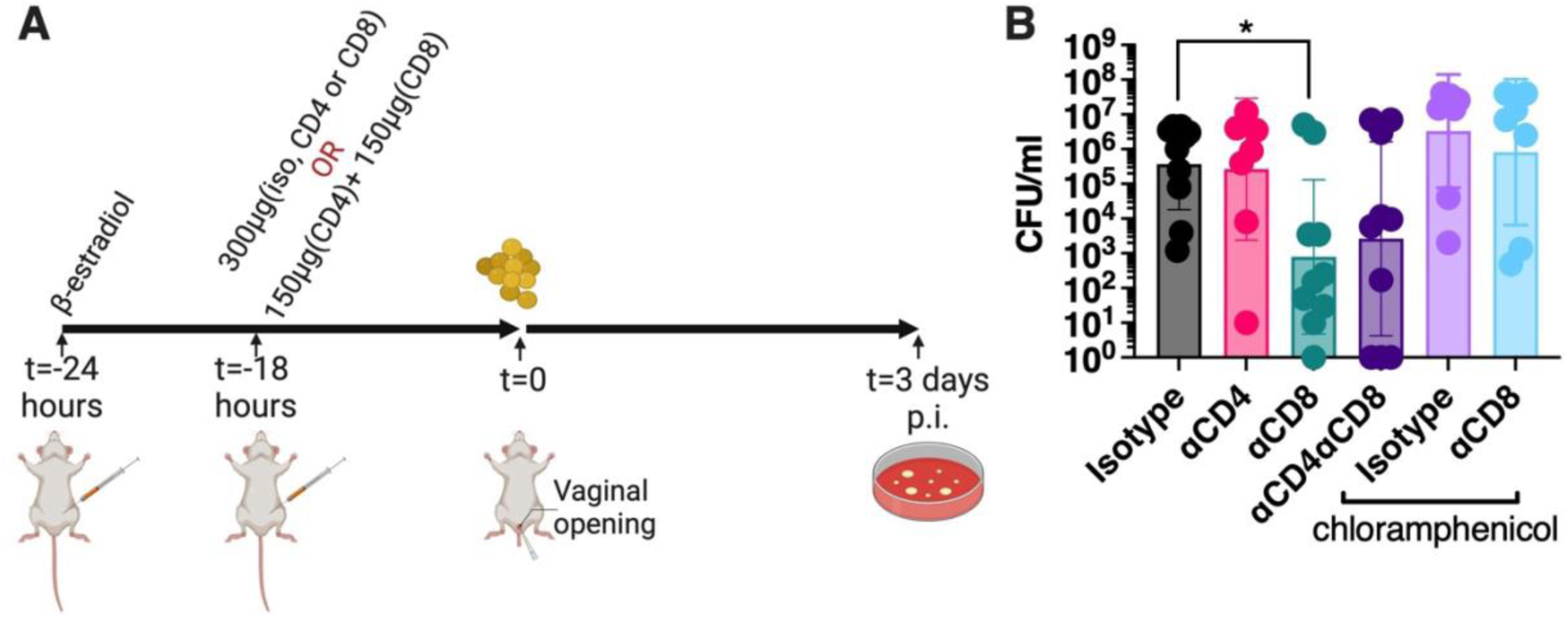
CD8^+^ T cells promote vaginal colonization by *S. aureus*. **(A)** Experimental T cell depletion timeline. **(B)** Mice were supplied with water *ad libitum*, either without or with chloramphenicol. The animals were all intraperitoneally injected with 300μg of an isotype control antibody, 300μg of anti-CD4 or anti-CD8 depleting antibodies, or a mix of 300μg (150μg of CD4-depleting antibodies and 150μg of CD8-depleting antibodies), 18 hours prior to vaginal inoculation with ∼10^6^ wild-type *S. aureus* MN8 in β-estradiol synchronized mice. Mice were sacrificed at day 3 and data are presented as the geometric mean ± SD. Each dot represents an individual mouse. Data were evaluated for significant differences using the Kruskal-Wallis test with multiple comparisons (*, *p*<0.05).

Lastly, to assess if CD8^+^ T cells were central to *S. aureus* colonization within the microbiota, CD8-depleting antibodies, or an isotype control, were injected in chloramphenicol treated animals. In contrast to mice with their endogenous microbes, the *S. aureus* burden did not decrease when CD8^+^ T cells were depleted (**Fig 4B**). These collective data suggest that superantigenic activation of CD8^+^ T cells was key for the function of TSST-1 on an intact vaginal microbiota to provide an advantage to *S. aureus* for vaginal colonization.

## Discussion

Bacterial superantigens are a group of extremely potent immunostimulatory toxins with a well-recognized role in both staphylococcal and streptococcal TSS (3). Of these toxins, the TSST-1 superantigen is considered to be the sole determinant of mTSS (20); however, a biological function for TSST-1 that is beneficial in the life-cycle of *S. aureus* has yet to be elucidated (4). Furthermore, mTSS is a rare disorder and occurs under a very restricted set of conditions, yet given the extreme potency of superantigens, production of TSST-1 at low levels may have an unrecognized impact within the vaginal niche without provoking mTSS. Herein, we evaluated the functional consequences of TSST-1 within the vaginal environment using *S. aureus* MN8, a well-characterized TSST-1^+^ and prototypic mTSS strain in a mouse model susceptible to this SAg (**Fig 1C**). We demonstrate that TSST-1 production was directly linked to an advantage for *S. aureus* during vaginal colonization, that TSST-1 impacted colonization during diestrus, and that this activity relied on CD8^+^ T cells. Furthermore, the TSST-1-dependent colonization phenotype was lost by depleting the vaginal microbiota by antibiotic treatment, providing evidence that the inflammatory activity of TSST-1 functions as a novel inter-bacterial competition factor within the vaginal tract.

Within the fluctuating vaginal environment in human, *S. aureus* will be exposed to multiple colonization barriers including low pH, host and bacterial antimicrobial molecules, direct competition with the established microbiota, and the host immune response. Given the dynamic changes that occur within the vaginal environment, mice were first synchronized into proestrus by β-estradiol. The estrous cycle then occurred over ∼4 days which was confirmed by the analysis of sex hormones (**Fig 1G**) (15). During diestrus there is an influx of leukocytes within the vaginal lumen which could impact the observed changes in *S. aureus* burden in the presence or absence of TSST-1 (15, 21). Although percentages of T cell populations were not detectably altered between mice colonized with either the wild-type or Δ*tst* mutant, wild-type MN8 induced a rapid increase in the inflammatory response by ∼24 hours post-inoculation, that subsequently decreased to levels that were similar to the MN8 Δ*tst* strain by ∼ 72 hours (**Fig 3**). In the absence of this TSST-1-induced inflammatory response, *S. aureus* CFUs decreased by ∼100-fold by days 3-4, and subsequently rebounded by day 7. This inflammatory response may be reminiscent to what is also observed during more classical bacterial sexually transmitted infections caused by *Treponema pallidum*, *Neisseria gonorrheae* or *Chlamydia trachomatis* (22, 23). With these sexually transmitted infections, this inflammatory phenotype has been associated with an increased susceptibility to the human immunodeficiency virus (HIV) as CD4^+^ are the main target of the virus (24–26).

The human vaginal mucosa possesses several immune particularities including: 1) the absence of an associated lymphoid center which will lead to few functions supported directly at the mucosa, including both immune priming of naïve CD8^+^ T cells but also expansion of antigen-specific CD8^+^; 2) a larger proportion of T cells but with approximately equal proportions of CD4^+^ and CD8^+^ T cells; and 3) relative consistency in the number of immune cells through the menstrual cycle compared to fluctuations that occur in the uterine mucosa (27–29). Although some of these characteristics can vary between humans and mice (e.g. CD4/CD8 proportion and a consistent number of vaginal T cells), this murine model allows for the study of complex interactions that occur with the presence of TSST-1 within the vaginal mucosa. Bacterial superantigens activate both CD4^+^ and CD8^+^ T cells, and herein we demonstrate maintenance of *S. aureus* within the vaginal tract required CD8^+^ T cells, but not CD4^+^ T cells (**Fig 4B**). This phenotype was in contrast to experimental bacteremia studies where the SEB and SEC staphylococcal superantigens promoted liver abscess formation through a CD4-dependent and excessive IFNγ response (30). Alternatively and similar to the current study, *Streptococcus pyogenes* appeared to require CD8^+^ T cells for efficient nasal carriage (31), but conversely in the nares, staphylococcal superantigens actually decreased colonization levels by *S. aureus* (32). These studies provide a picture where superantigens have evolved niche-specific functions, that are most likely related to tissue specific immunity, such that the production of these toxins can favor the establishment of the bacterium through multiple but tissue-dependent mechanisms. Although the downstream mechanisms following activation of CD8^+^ cells by superantigen requires further investigation, the present work suggests that TSST-1 can manipulate this cellular response to compete against the microbial species in the murine vaginal tract.

*S. aureus* can colonize multiple sites, and each mucosal environment will present different environmental signals, which will subsequently lead to differential expression of virulence factors. In the human vaginal tract, a major signal that will repress TSST-1 expression is vaginal glucose levels, where TSST-1 is repressed specifically by CcpA (33). However, glucose levels are highly variable as they are released from the glycogen produced by the vaginal epithelium and these cells are both producing glycogen and exfoliated depending on estrogen levels (34–37). Depending of this fluctuation of glucose repression, TSST-1 will be produced at different times and different quantities during the menstrual cycle, not usually at levels needed to invoke mTSS, but still at concentrations able to manipulate the immune response (20) (**Fig. 2 and 3**). In the murine model of this study, 10 μg TSST-1 inoculated in proestrus is suggested to be similar to the dose secreted by MN8 producing TSST-1 and is sufficient to provoke changes in *S. aureus* colonization **(Fig. 1F)**. Through this process, the production of TSST-1 appears to favor *S. aureus* growth and colonization within the vaginal environment.

## Material and methods

### Bacterial strains and growth conditions

Bacterial strains used for this study are listed in **Table 1**. *S. aureus* strains were grown aerobically at 37°C in tryptic soy broth (TSB) (Difco) or brain heart infusion (BHI) broth with shaking, or on TSB plate at 37°C with antibiotics as needed. *Escherichia coli* XL1 Blue was used as a cloning host and *E. coli* BL21(DE3) was used for expression of recombinant TSST-1 and its inactive form (TSST-1_S72A_). *E. coli* strains were grown in Luria Bertani (LB) broth with appropriate antibiotics at 37°C with aeration or grown aerobically in TSB at 37°C, 250 rpm, or on TSA (tryptic soy agar) plates with appropriated antibiotics.

To generate a markerless deletion of the *tst* gene in *S. aureus* MN8, ∼1,029-bp of DNA immediately upstream of *tst* was PCR amplified using the primers *tst*_Upstream_attB1_F (GGGGACAAGTTTGTACAAAAAAGCAGGCTTTTATACATACACCTAAT ATGTTT) and *tst* Upstream R (TTTATTCATTTTTAATTCTCCTTCAT), and a 1,009-bp of DNA immediately downstream of *tst* was amplified using the primers *tst*_Downstream_F (ATTAATTAATTTACCACTTTTTCTG*) and *tst*_Downstream_attB2 (GGGGACCACTTTG TACAAGAAAGCTGGGTGTTGATAGATGATGAAATAAATAC). These products were cloned into the pKOR1 integration vector (38) using the Gateway BP Clonase II system (Life Technologies). This plasmid was passaged through *E. coli* SA30B to methylate the DNA (39) and the *tst* deletion was introduced into the *S. aureus* MN8 chromosome as described (40). The *tst* complementation vector was constructed by amplifying *tst* using the primers *tst*_comp_F_KpnI (ACACGGTACCGCTCCCTATGTAACAAACACTTTT) and *tst*_comp_R_EcoRI (CCGAATTCAAAGATAAAAGGGAGAACGCTTA), and cloned into the complementation plasmid pCM29 using *Kpn*I and *Eco*RI. This plasmid confirmed by whole-plasmid sequencing (Plasmidsaurus) and along with empty pCM29, were separately transformed into *S. aureus* strains by electroporation after the methylation step into *E. coli* SA30B. Growth of these strains were assessed in TSB broth with agitation for 18 hours in Biotek Synergy H4 multimode plate reader.

### Mice

Female BALB/c between 7- and 9-weeks old were used for all *in vivo* experiments and were purchased from either Jackson Laboratory (stock 000651) or Charles River Laboratory (stock 028). Animals were housed during experiments without exceeding 4 animals per cage. Mice were provided water and food *ad libitum* and appropriate enrichment was added to all cages. All experiments were in accordance with the Canadian Council on Animal Care Guide to the Care and Use of Experimental Animals, and the animal protocol was approved by the Animal Use Subcommittee at the University of Western Ontario (Protocol #2020-061).

### Murine splenocyte analysis

The ability of murine T cells to respond to TSST-1 was tested through production of IL-2 from mouse splenocytes (14). Briefly, spleens were homogenized and red blood cells were lysed with ammonium-chloride-potassium (ACK) buffer. The remaining cells were resuspended in RPMI supplemented by 10% fetal bovine serum (Wisent), 2mM glutamine (Wisent), 1mM sodium pyruvate (Gibco), 100µM non-essential amino acids (Gibco), 25mM HEPES pH 7.2 (Gibco), 100ug/ml streptomycin (Gibco), 100U/ml penicillin (Gibco) and 2µg/ml polymyxin B (Gibco). Splenocytes were seeded in 96-wells plate at a final concentration of 1 x 10^6^ cells/ml and concentrations of TSST-1 from 1pg/ml to 10ug/ml were added to cells and incubated for 18 h at 37°C with 5% CO_2_. Supernatants issued from TSST-1 challenge were assessed for IL-2 concentration by enzyme-linked immunosorbent assay (ELISA) according to manufacturer’s protocol (Invitrogen). The plates were read at 450nm and 570nm in Biotek Synergy H4 multimode plate reader.

### Murine vaginal colonization model

Twenty-four hours prior to vaginal inoculation, 50 μg of β-estradiol resuspended in commercial canola oil was injected intraperitoneally. On the day of inoculation, strains were subcultured in fresh TSB, resuspended in HBSS, and 2.5 (± 2.5) x 10^6^ bacteria were administered intravaginally in a 10 μl dose. Animals were sacrificed at indicated time points post-inoculation and the lower reproductive tract was excised, homogenized and bacterial counts were assessed after incubation on Mannitol-Salt Agar (MSA – Fisher Scientific) plates.

### Hormones and cytokines/chemokines analyses

At various time points post-inoculation, serum (6, 24, 48 and 72 hours) and vaginal homogenates (2, 8, 12, 24, 48 and 72 hours) were collected. Vaginal homogenates were obtained by homogenization in HBSS supplemented with the complete protease inhibitor mixture (Roche). Vaginal samples were analyzed using Steroid/Thyroid Hormone 6-plex Discovery Assay (Eve Technologies) or Mouse Cytokine/Chemokine 32-Plex Discovery Assay (Eve Technologies). Serum samples were obtained by blood draw at the posterior vena cava after coating the needle with heparin. Serum was separated from erythrocytes and leukocytes through centrifugation. Samples were filter-sterilized and were analyzed using Mouse Cytokine/Chemokine 32-Plex Discovery Assay (Eve Technologies). Both serum and homogenates results are presented as the percentage of the highest averaged concentration of a cytokine. Significance for each cytokine/chemokine at each timepoint was assessed in between the 2 groups using Mann Whitney U test.

### Microbiota depletion

For depleting the vaginal microbiota, the same procedure for the vaginal colonization model was followed with sacrifice at 3 days post-inoculation although at 2 days prior to bacterial inoculation, chloramphenicol (2mg/L) was added to drinking water and was refreshed every other day (t=0 and 2 days p.i.) until sacrifice. Vaginas were homogenized and incubated on MSA supplemented with chloramphenicol for *S. aureus* CFU determination. Additional plates issued from the same homogenates were incubated to confirm microbiota depletion. MSA agar without antibiotic (other staphylococci), blood plates (differentiation in between various species – Hardy Diagnostics), and Enterococcus selective media (detection of *Enterococcus* species - Sigma Aldrich) were incubated at 37°C in atmospheric level of oxygenation for 24 hours. MRS agar plates (detection of presumed lactobacilli) were incubated in an anaerobic jar with GasPak EZ (Fisher Scientific) at 37°C for 24 hours.

### Vaginal inoculation of TSST-1 and 16S rRNA gene sequencing

Recombinant TSST-1 and an attenuated version of TSST-1 carrying an S72A point mutation (TSST-1_S72A_) were produced as described (41). Briefly, the 2 variants were expressed with a His-tag in *E. coli* BL21(DE3) and purified by nickel column chromatography. A 10 μg dose of TSST-1 or the inactive mutant TSST-1_S72A_ was then inoculated alone in mice for 3 days and vaginal tract were collected and frozen for 16S sequencing. The use of TSST-1 and its variant in experimental settings is approved by University of Western Ontario Biosafety Committee (BIO-UWO-0155).

For microbiome analysis, tissue sections from the internal vaginal tract were extracted with the Qiagen DNeasy Powersoil Pro kit as previously described (42). Extracted DNA was stored at −20°C until PCR amplification. PCR amplification was completed using the Bakt_341F (5′-CCTACGGGNGGCWGCAG-3′) and Bakt_805R (5′-GACTACHVGGGTATCTAATCC-3′) universal primer set, which targets the V3–V4 variable region of the 16S rRNA gene enabling species-level classification of bacteria. Sequencing was carried out at the London Regional Genomics Center (http://www.lrgc.ca). First, a pippin preparation was used to size-select the 16S amplicons and exclude host amplicons. Amplicons were then quantified using pico green and pooled at equimolar concentrations before cleanup using QIAquick (Qiagen). Amplicons were sequenced with 2 x 300 bp paired-end chemistry on the Illumina MiSeq using the 600-cycle MiSeq v.3 Reagent Kit. Following sequencing, paired reads were exported as fastq files (uploaded to NCBI Sequence Read Archive, BioProject ID PRJNA1117910). Demultiplexed reads were then quality filtered, trimmed, denoised, and merged following the DADA2pipeline (version 1.26.0) in R (version 4.2.1) (43, 44). The SILVA Database (version 138) was utilized in assigning taxonomy to the amplicon sequence variants (SVs). SVs were filtered such that those not comprising at least 0.1% of the relative abundance in any sample were removed. Samples were filtered by read count: only samples with a read count of at least 1000 were included. For samples with >1000 reads, read count, taxonomic, and sequence information are in Supplementary Data. Alpha diversity metrics were determined with the R package phyloseq (version 1.42.0) (45).

### Flow cytometry analysis of vaginal cells

Vaginal tracts were extracted from mice cells were isolated and washed as previously described (46). Following isolation, vaginal cells were first stained for cell viability using Fixable Viability Dye eFluor506 (Thermo-Fisher) and then subsequently stained with anti-CD45-BV421 (clone 30-F11, BioLegend), anti-CD4-PE-Cy5 (clone RM4-5, Thermo-Fisher), anti-CD8a-PE-Cy7 (clone 53-6,7, BioLegend), anti-CD19-BV711 (clone 1D3, BD Biosciences) and anti-F4/80-A700 (clone RB6-8C5, BioLegend). Events were acquired using LSR II (BD Biosciences) and analyzed using FlowJo v10.7.1 (TreeStar). The gating strategy is presented in **Figure S2**. Results are presented as either the cells counts (CD45^+^ cells) or the percentage of CD45^+^ cells (CD4^+^ and CD8^+^ cells) for each animal from three biological replicates.

### CD4^+^ and CD8^+^ T cells depletion

The T cell depletion strategy was adapted from Tuffs *et al*, 2022 (30). Twenty-four hours prior to inoculation, 50 μg of β-estradiol is intraperitoneally injected, then 18 hours prior to bacterial inoculation 300 μg of depleting antibodies either anti-CD4 (clone GK1.5, BioXCell), anti-CD8 (clone YTS169.4, BioXCell) or a Rat IgG2b isotype control (clone LTF-2, BioXCell) was intraperitoneally injected. For double depletion, both anti-CD4 and anti-CD8 were injected at 150 μg each. *S. aureus* MN8 wild type was inoculated at ∼10^6^ bacteria per dose intravaginally, and mice were sacrificed at 3 days post-inoculation. Vaginas were homogenized and incubated on MSA supplemented with chloramphenicol for bacterial burden. Cell depletions were confirmed by flow cytometry from CD45^+^ vaginal homogenates extracted as above (**Fig. S4**).

### Statistical analysis

Statistical analyses were performed in R or using GraphPad Prism 9 and a *p* value equal or lower than 0.05 was considered statistically significant. For bacterial burden CFU, nonparametric Mann-Whitney U test or Kruskal-Wallis test with an uncorrected Dunn’s test for multiple comparisons were performed. For microbiota differential abundance comparisons, the R package ALDEx2 was used to determine effect size, for which ≥|0.5| was considered statistically significant (47).

## Acknowledgements

This work was supported by funding from the Canadian Institutes of Health Research (CIHR) Grant PJT-166050 to JKM and JB acknowledges funding from the Weston Family Foundation. KFA acknowledges support from a Research Scholars Award from the American Urological Association.

